# Predicting genes associated with RNA methylation pathways using machine learning

**DOI:** 10.1101/2021.12.10.472055

**Authors:** Georgia Tsagkogeorga, Helena Santos-Rosa, Andrej Alendar, Dan Leggate, Oliver Rausch, Tony Kouzarides, Hendrik Weisser, Namshik Han

## Abstract

RNA methylation plays an important role in functional regulation of RNAs, and has thus attracted an increasing interest in biology and drug discovery. Here, we collected and collated transcriptomic, proteomic, structural and physical interaction data from the Harmonizome database, and applied supervised machine learning to predict novel genes associated with RNA methylation pathways in human. We selected five types of classifiers, which we trained and evaluated using cross-validation on multiple training sets. The best models reached 88% accuracy based on cross-validation, and an average 91% accuracy on the test set. Using protein-protein interaction data, we propose six molecular sub-networks linking model predictions to previously known RNA methylation genes, with roles in mRNA methylation, tRNA processing, rRNA processing, but also protein and chromatin modifications. Our study exemplifies how access to large omics datasets joined by machine learning methods can be used to predict gene function.

## INTRODUCTION

RNA modifications have been known since the 1960s, when the sequencing of the first transfer RNA (tRNA) from yeast revealed 10 chemically modified ribonucleosides, including pseudouridine (Ψ)^1^. Since then, the number of identified modifications has grown to over 150, found on both coding and non-coding RNAs across all three kingdoms of life^2^. Technological advances in the field have established that RNA modifications are widespread, reversible and dynamically regulated^1^. Methylation is the most abundant type, with methyl-groups decorating multiple RNA species, such as messenger RNA (mRNA), ribosomal RNA (rRNA) and tRNA, at different nucleosides and positions. So far, N6-methyladenosine (m^6^A) is the most studied modification, commonly detected in mRNA, rRNA, long intergenic non-coding RNA (lincRNA), primary microRNA (pri-miRNA), and small nuclear RNAs (snRNA). Other methyl-marks include 5-methylcytosine (m^5^C), N1-methyladenosine (m^1^A), 7-methylguanosine (m^7^G), 2’-O-dimethyladenosine (m^6^Am) and 5-hydroxymethylcytosine (hm^5^C)^3–5^.

Deposition of methyl-marks on RNA is catalysed by writer enzymes, known as RNA methyltransferases. To date, there are 57 RNA methyltransferases identified in the human genome. Of these, five methylate mRNAs, six small RNAs, 14 rRNAs, and 22 tRNAs, whereas 12 remain with unknown substrates^6^. Most enzymes use S-adenosyl-methionine (SAM) as a methyl donor to the RNA substrate, while many also recruit accessory proteins, which are often essential for substrate binding, localization, and stability. The most well-studied examples of RNA methylation writers are by far the complex METTL3-METTL14 complex responsible for the deposition of m^6^A, followed by a NOL1/NOP2/Sun (NSUN) domain-containing family of tRNA-modifying enzymes depositing m^5^C on tRNAs^7^.

Dynamic regulation of RNAs via chemical modifications has recently attracted a rising interest in RNA modifying enzymes as new potential therapeutic targets^8^. This is because multiple lines of evidence suggest that RNA methylation plays a far more important role in cell functioning than previously thought. In line with this, several studies have shown that RNA methylation is a key modulator of transcript stability, gene expression, splicing and translation efficiency^9–11^. Furthermore, a growing body of data has demonstrated that changes in RNA methylation processes can be linked to a range of cancers, neurological disorders and various other diseases^12^. Surprisingly, despite this critical role in cellular homeostasis and disease, RNA methylation pathways in general remain understudied^7^. Our current understanding of RNA modifications is also highly fragmentary, with an estimated 20% or more of RNA modifying enzymes still remaining unknown or unidentified^13^.

Conventional approaches for studying novel gene functions include a range of labour-intensive wet-lab techniques, including mutagenesis, gene disruption or gene depletion (knocking-down/-out) for characterising gene-specific phenotypic effects, and chromatography and mass spectrometry for identifying molecular interactions. However, over the last two decades, access to large-scale omics data has enabled the use of “dry” computational methods for understanding biological functions. A wide array of bioinformatic tools have been developed under the umbrella of functional genomics, ranging from methods used to identify homologous genes with similar functionalities across species to genome-wide screens for specific sequence motifs and functional domains. Today, machine learning techniques are emerging as a powerful approach to harness the increasing wealth of large-scale biological data, allowing the discovery of hidden patterns and more reliable statistical predictions^14^.

Here, we aimed to better understand the molecular pathways involved in RNA methylation in human using machine learning. To this end, we used publicly available human transcriptomic, proteomic, structural and protein-protein interaction data^15^ and built a large machine learning dataset for supervised binary classification. We trained and evaluated five ensembles of predictive models: Logistic Regression (LR), Gaussian Naïve Bayes (GNB), Support Vector Machine (SVM), Random Forest (RF) and Gradient Boosting (GB) models. We employed the best models to predict genes functionally associated with RNA methylation pathways in the human genome.

## RESULTS AND DISCUSSION

### Data engineering and feature selection

Mining functional annotation databases in conjunction with extensive literature searches allowed us to identify 92 proteins involved in RNA methylation (Table 1). These were either methyl-writers (known RNA methyltransferases6 and their partner proteins in protein complexes), or enzymes previously annotated as putative RNA methyltransferases (see Methods). Genes encoding for these proteins constituted our positive class (Class 1) in machine learning analyses. To frame our predictive modelling as a binary classification problem, we assembled multiple stratified training and test datasets by randomly sampling a number of genes equal to our positive set from the remaining genome, ensuring that all genes of our initial dataset were sampled exactly once (Figure 1). Our rationale was that this would allow machine learning models to be trained and tested across a diverse range of other gene functions, instead of just choosing one function for the negative set. In addition, this approach alleviates any putative bias that may arise from sampling a single negative set of genes from the human genome.

**Figure 1.**
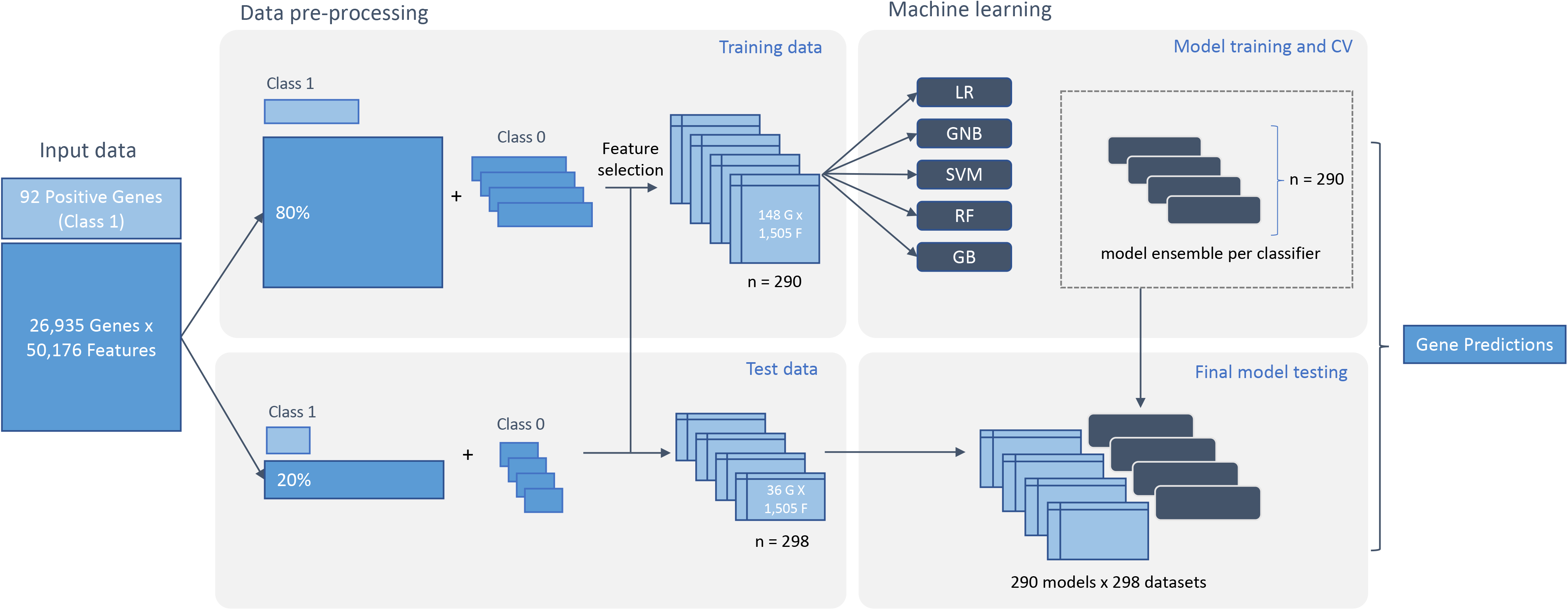
Schematic representation of the analysis workflow. Previously known RNA methylation genes were used as positive samples (Class 1) and split into two sets comprising 80% of the data for training and 20% kept unseen for model testing. An analogous 80/20 split was performed for the remaining genes of the human genome, which were further divided into sets of equal size to the positive samples and used as negative samples (Class 0) to generate stratified sets for training and testing. Following feature pre-filtering, five types of machine learning models for binary classification - Logistic Regression (LR), Gaussian Naïve Bayes (GNB), Support Vector Machine (SVM), Random Forest (RF) and Gradient Boosting (GB) - were trained on each of the training sets resulting in a classifier ensemble. Each model from the classifier ensemble was evaluated on each of the test datasets and overall performance was calculated by averaging results of all models across test sets. The best-performing ensemble was used to make predictions for the whole genome.

**Table 1.**
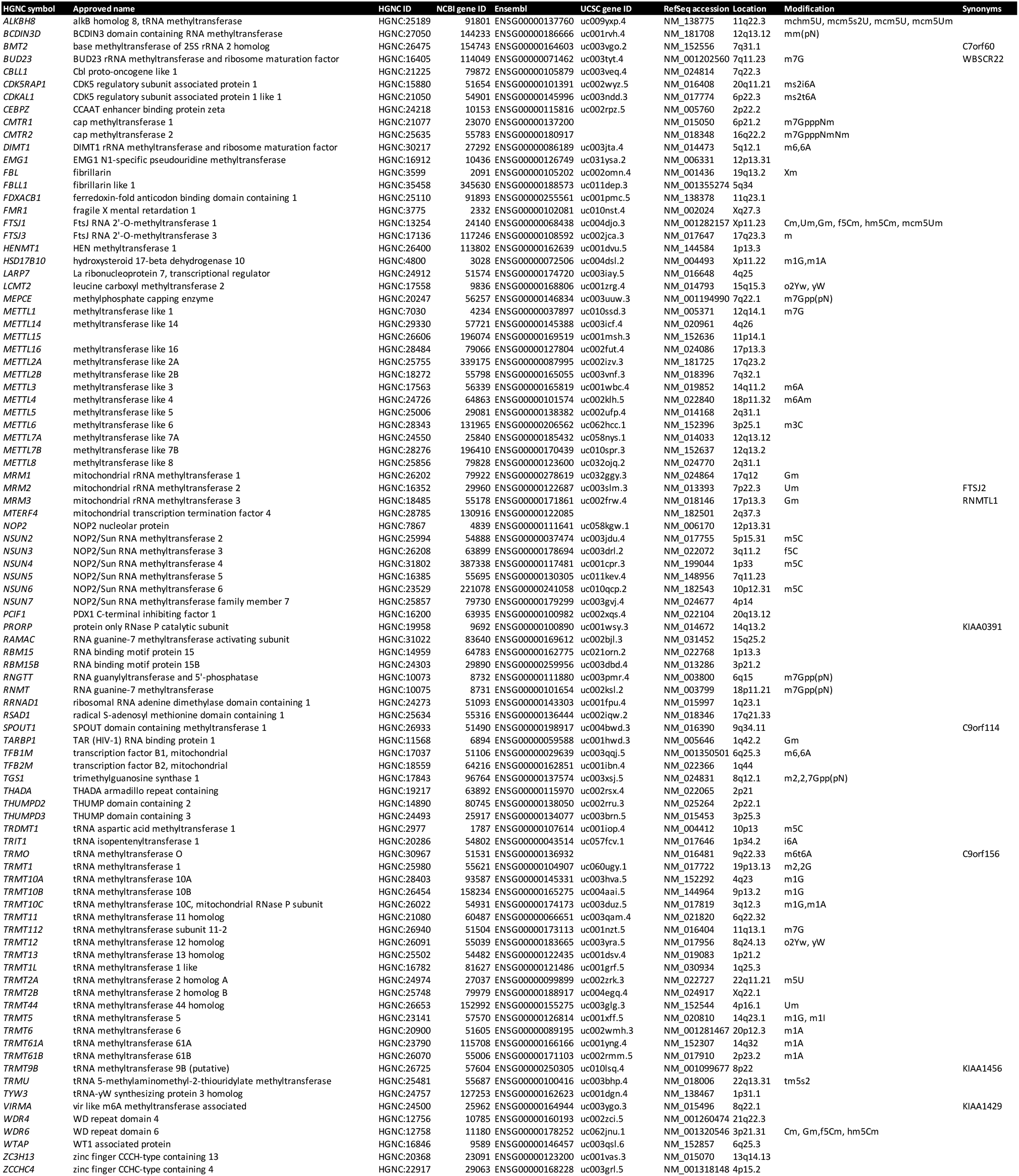
Known RNA methyltransferases and related proteins used as positive set (Class 1).

We initially pooled 50,176 features collected from publicly available and previously curated transcriptomic, proteomic, functional annotation, structural and physical interaction datasets (Table 2). To identify features that were informative for classification and thereby useful for predicting genes associated with RNA methylation, we performed feature selection prior to model training, followed by feature ranking after training and cross-validation. To reduce the feature-to-sample ratio, first we eliminated features with excessive missing data in the training dataset. Second, we removed features with low variance, which resulted in a drastic dimensionality reduction to 1,505 features for the final dataset. Selected features used for classification were drawn from BioGPS^16^ (35), Gene Ontology^17^ (GO: 59), GTEx^18^ (1,114), Human Protein Atlas^19^ (HPA: 107), InterPro (1), Pathway Commons (PathCommons: 150) and TISSUES^20^ (40) datasets.

**Table 2.** Gene-feature omics datasets used in machine learning analyses (source Harmonizome).

| Dataset | Description | Measurement | Association | Category | Resource | Genes | Attributes | Associations |
| --- | --- | --- | --- | --- | --- | --- | --- | --- |
| BioGPS Human Cell Type and Tissue Gene Expression Profiles Dataset | mRNA expression prof | Gene expression by microa | Gene-cell type or tissue associations by differential expression of | Transcriptomics | BioGPS | 16379 | 84 cell type or tissues | 205445 gene-cell type or tissue associations |
| GTEx Tissue Gene Expression Profiles Dataset | mRNA expression prof | Gene expression by RNA-si | Gene-tissue associations by differential expression of gene across | Transcriptomics | Genotype Tissue Expression | 25557 | 29 tissues | 112583 gene-tissue associations |
| GTEx Tissue Sample Gene Expression Profiles Dataset | mRNA expression prof | Gene expression by RNA-si | Gene-tissue sample associations by differential expression of gene across | Transcriptomics | Genotype Tissue Expression | 19247 | 2918 tissue samples | 8421199 gene-tissue sample associations |
| HPA Cell Line Gene Expression Profiles Dataset | mRNA expression prof | Gene expression by RNA-si | Gene-cell line associations by differential expression of gene across | Transcriptomics | Human Protein Atlas | 15372 | 43 cell lines | 102943 gene-cell line associations |
| HPA Tissue Gene Expression Profiles Dataset | mRNA expression prof | Gene expression by RNA-si | Gene-tissue associations by differential expression of gene across | Transcriptomics | Human Protein Atlas | 17423 | 31 tissues | 81082 gene-tissue associations |
| HPA Tissue Sample Gene Expression Profiles Dataset | mRNA expression prof | Gene expression by RNA-si | Gene-tissue sample associations by differential expression of gene across | Transcriptomics | Human Protein Atlas | 16657 | 121 tissue samples | 303267 gene-tissue sample associations |
| GO Biological Process Annotations Dataset | Curated annotations of | Association by literature c | Gene-biological process associations from curated gene annotations | Structural or functional annotations | Gene Ontology | 15717 | 13212 biological processes | 969303 gene-biological process associations |
| GO Molecular Function Annotations Dataset | Curated annotations of | Association by literature c | Gene-molecular function associations from curated gene annotations | Structural or functional annotations | Gene Ontology | 15777 | 4162 molecular functions | 223181 gene-molecular function associations |
| InterPro Predicted Protein Domain Annotations Dataset | Protein domains predicted | Association by computational | Protein-protein domain associations by sequence similarity to domain | Structural or functional annotations | InterPro | 18002 | 11015 protein domains | 62614 gene-protein domain associations |
| KEGG Pathways Dataset | Sets of proteins participating | Association by literature c | Protein-pathway associations from curated pathways | Structural or functional annotations | Kyoto Encyclopedia of Genes | 3947 | 200 pathways | 9324 gene-pathway associations |
| Reactome Pathways Dataset | Sets of proteins participating | Association by literature c | Protein-pathway associations from curated pathways | Structural or functional annotations | Reactome | 7535 | 1638 pathways | 83680 gene-pathway associations |
| TISSUES Curated Tissue Protein Expression Evidence Scores Dataset | Protein tissue expression | Association by literature c | Protein-tissue associations by integrating evidence from manual | Proteomics | TISSUES | 16215 | 643 tissues | 357442 gene-tissue associations |
| HPA Tissue Protein Expression Profiles Dataset | Semiquantitative protein | Protein expression by immun | Protein-tissue associations by differential expression of protein across | Proteomics | Human Protein Atlas | 15704 | 44 tissues | 138576 gene-tissue associations |
| Hub Proteins Protein-Protein Interactions Dataset | Sets of proteins interacting | Association by data aggregation | Protein-hub protein associations from aggregated protein-protein | Physical interactions | Hub Proteins | 9362 | 289 hub proteins | 58320 gene-hub protein association |
| Pathway Commons Protein-Protein Interactions Dataset | Protein-protein interactions | Association by data aggregation | Protein-protein associations from low-throughput or high-throughput | Physical interactions | Pathway Commons | 15747 | 15747 interacting proteins | 3527164 gene-interacting protein associations |

During model training and cross-validation, we computed feature importance by using the GB importance measure as averaged across all training sets. The 50 most informative features and their relative importance in classification are shown in Figure 2. The features with the highest importance for the full feature set were mainly GO terms, such as GO:0032259, GO:0016740, GO:0003723, GO:0008168 and GO:0016070, all corresponding to methylation, transferase/methyltransferase activity and RNA metabolic processes. Equally, the InterPro domain IPR029063, which represents the S-adenosyl-L-methionine-dependent methyltransferase superfamily was ranked among the top 50 most informative features (Figure 2A). Although anticipated, the fact that the classifiers seemed to rely on RNA and methylation-related annotation features provides support that the models learn to classify genes with a strong link to RNA methylation processes.

**Figure 2.**
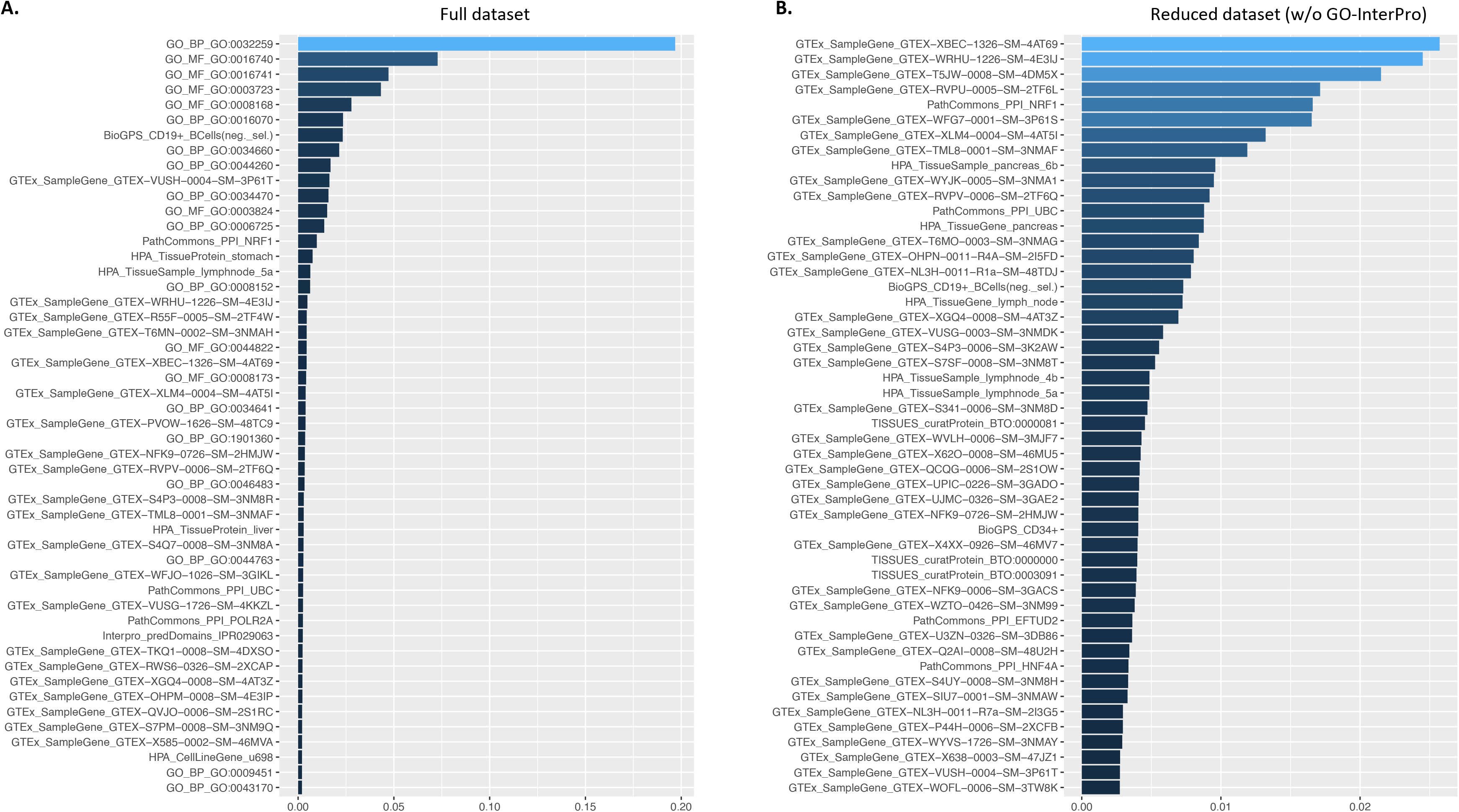
Feature importance. Top 50 most informative features ranked by their relative importance in predictive modelling based on the **A.** full and **B**. reduced feature sets.

Although GO annotations are informative, they may equally bias gene prediction towards pre-existing functional annotations. We assembled thus a second feature set of reduced dimensionality, by excluding GO and InterPro data types. When classifiers were trained on this reduced feature set, the most informative types of features were mainly GTEx expression profiles (Figure 2B). The GTEx project aims to provide a comprehensive public resource of tissue-specific gene expression and regulation, so far including samples from 54 non-diseased tissues across nearly 1000 individuals^18^. Tissue sample expression data as integrated in Harmonizome and thus sampled here, consist of one-hot-encoded sets of genes with high or low expression in each tissue sample relative to other tissue samples from the GTEx tissue expression profiles dataset.

A possible interpretation of the high ranking of such GTEx expression profile features is that under specific biological conditions, i.e., in certain tissues, RNA methylation genes tend to be collectively down- or up-regulated as compared to other processes. Alternatively, a high ranking of GTEx features may be due to the high proportion of GTEx features in the feature set and noise originating from the high dimensionality of the training dataset with respect to the feature-to-sample ratio. To investigate this further, we calculated the relative frequency of GTEx features in the top hundred most informative features across models from all training sets (Table 3). Notably, certain samples taken from the areas of blood, heart, pancreas, and brain were retrieved as informative by more than a hundred models.

**Table 3.**
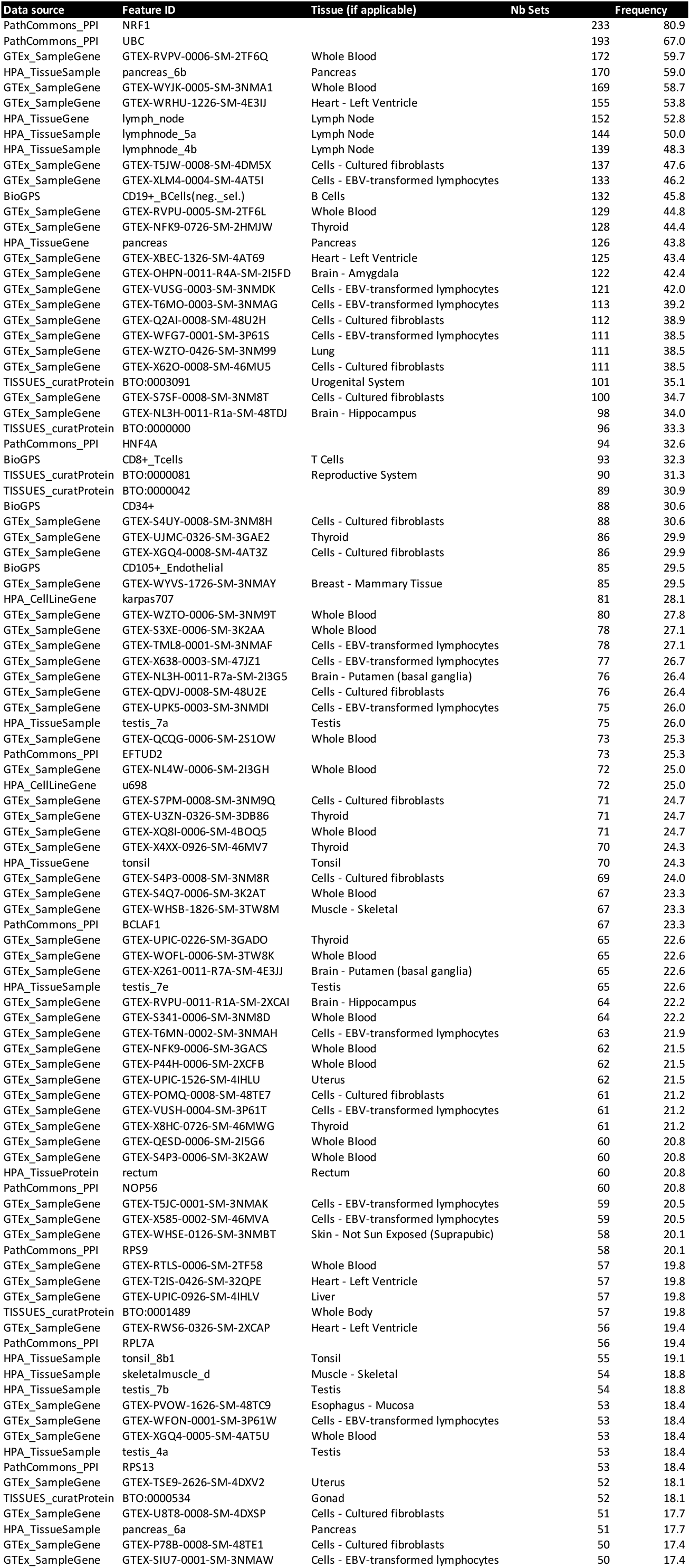
Highly informative features based on models trained on the reduced feature set, and their frequency in the top100 features across all models of the classifier ensemble.

### Model performance

We selected five machine learning classifiers (LR, GNB, SVM, RF and GB) and trained each on training sets from the full and the reduced feature set, creating an ensemble of models per classifier and feature set. To evaluate model performance, we used 10-fold cross validation and standard performance quantification metrics, i.e., accuracy, precision, recall, F1 score, and Area Under the Curve of the Receiver Operating Characteristic (AUCROC). Overall, all five model ensembles showed very similar performance based on cross-validation (Table 4). Among classifiers trained using the full feature set, GB and RF models showed the highest average accuracy at 0.875 and 0.870, respectively, as well as a similarly high average precision of 0.895 and 0.870, respectively. The GB ensemble followed by that the RF models also yielded the highest AUROC score, with an average AUC estimated at 0.938 and 0.937, respectively.

**Table 4.**
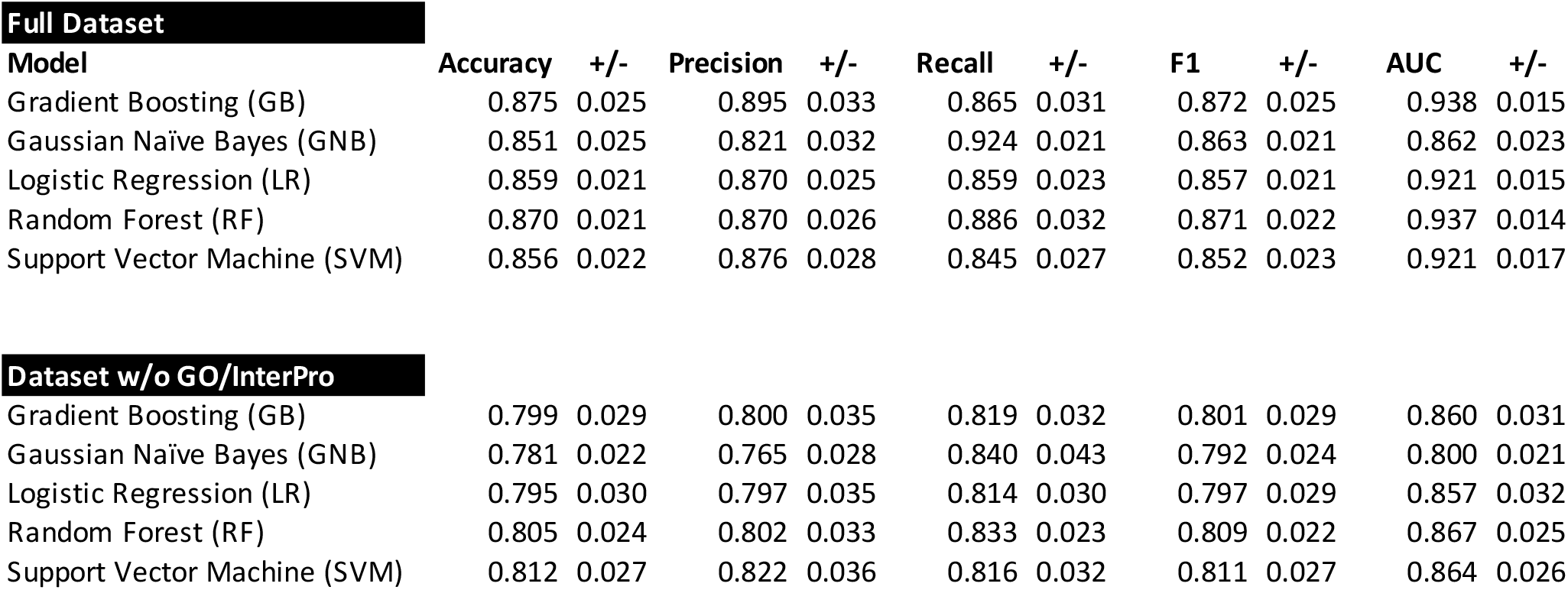
Model performance based on 10-fold cross-validation.

| Model | Accuracy | +/- | Precision | +/- | Recall | +/- | F1 | +/- | AUC | +/- |
| --- | --- | --- | --- | --- | --- | --- | --- | --- | --- | --- |
| Gradient Boosting (GB) | 0.875 | 0.025 | 0.895 | 0.033 | 0.865 | 0.031 | 0.872 | 0.025 | 0.938 | 0.015 |
| Gaussian Naïve Bayes (GNB) | 0.851 | 0.025 | 0.821 | 0.032 | 0.924 | 0.021 | 0.863 | 0.021 | 0.862 | 0.023 |
| Logistic Regression (LR) | 0.859 | 0.021 | 0.870 | 0.025 | 0.859 | 0.023 | 0.857 | 0.021 | 0.921 | 0.015 |
| Random Forest (RF) | 0.870 | 0.021 | 0.870 | 0.026 | 0.886 | 0.032 | 0.871 | 0.022 | 0.937 | 0.014 |
| Support Vector Machine (SVM) | 0.856 | 0.022 | 0.876 | 0.028 | 0.845 | 0.027 | 0.852 | 0.023 | 0.921 | 0.017 |
| **Dataset w/o GO/InterPro** |  |  |  |  |  |  |  |  |  |  |
| Gradient Boosting (GB) | 0.799 | 0.029 | 0.800 | 0.035 | 0.819 | 0.032 | 0.801 | 0.029 | 0.860 | 0.031 |
| Gaussian Naïve Bayes (GNB) | 0.781 | 0.022 | 0.765 | 0.028 | 0.840 | 0.043 | 0.792 | 0.024 | 0.800 | 0.021 |
| Logistic Regression (LR) | 0.795 | 0.030 | 0.797 | 0.035 | 0.814 | 0.030 | 0.797 | 0.029 | 0.857 | 0.032 |
| Random Forest (RF) | 0.805 | 0.024 | 0.802 | 0.033 | 0.833 | 0.023 | 0.809 | 0.022 | 0.867 | 0.025 |
| Support Vector Machine (SVM) | 0.812 | 0.027 | 0.822 | 0.036 | 0.816 | 0.032 | 0.811 | 0.027 | 0.864 | 0.026 |

The performance of the five classifiers for the reduced feature set without GO/InterPro annotations was diminished compared to the full dataset (Table 4). The model ensembles of SVM and RF outperformed the remaining three ensembles across almost all metrics. SVM models performed the best on the reduced feature set based on cross-validation, with an average prediction accuracy of 0.812, precision of 0.822 and AUROC of 0.864.

Based on the above results, we selected the best model ensembles to apply on previously unseen test data: GB for the full feature set and SVM for the reduced feature set. Accuracy, precision, recall and AUCROC for the test datasets were calculated by averaging the values obtained for each model in an ensemble. For the ensemble of GB models using the full feature set, the average test set accuracy was 0.905, precision 0.897 and recall 0.923 (Figure 3A). The average test set accuracy, precision and recall for SVM models trained on the reduced feature set were 0.830, 0.820 and 0.857, respectively (Figure 3). The average AUCROC was 0.973 for the GB model ensemble, and 0.899 for the SVM ensemble.

**Figure 3.**
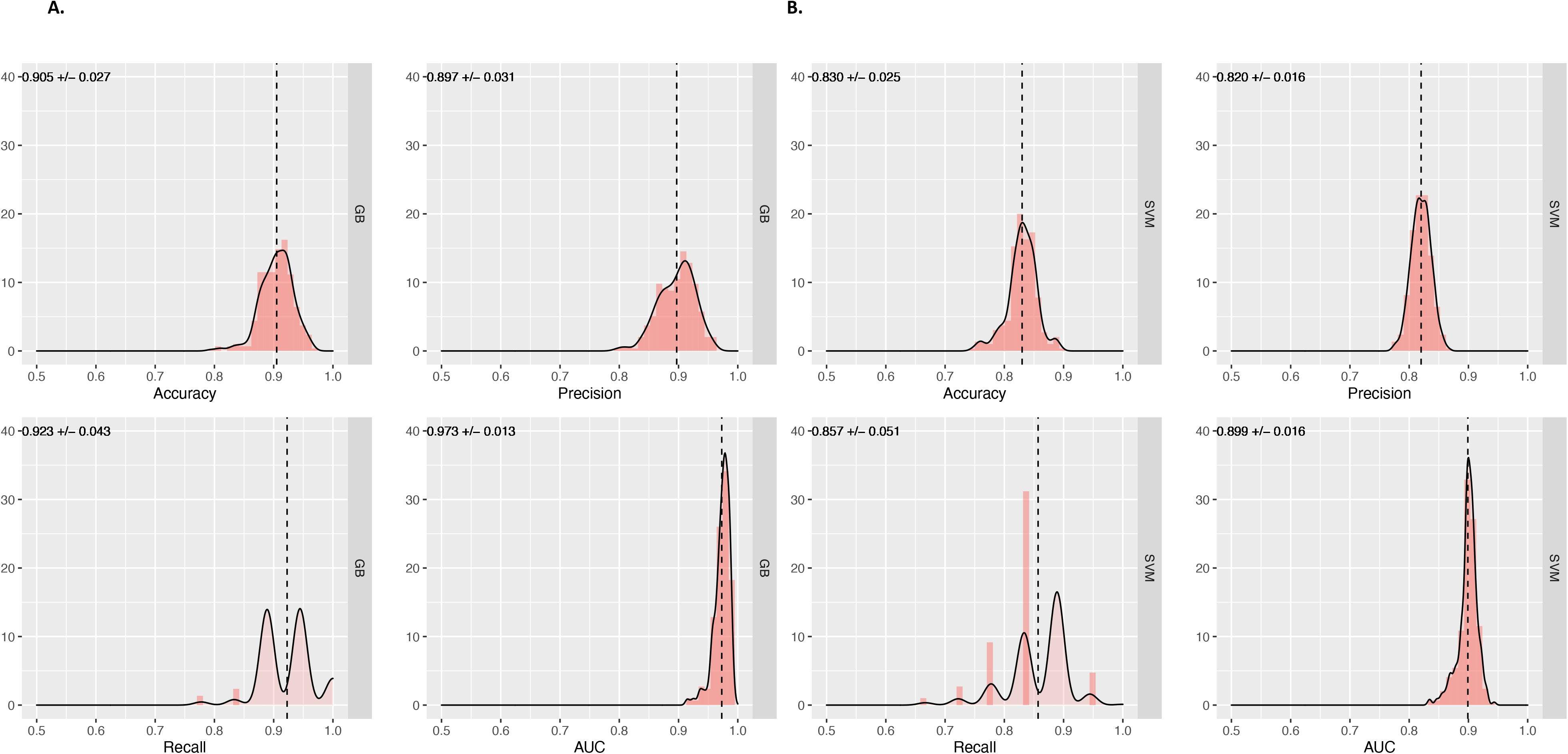
Model performance based on test data. Accuracy, precision, recall and AUC score distributions as estimated across test datasets for the best model ensembles: **A.** GB models for the full feature set; and **B.** SVM models for the reduced feature set.

### Model predictions and *in silico* validation

#### What do the models predict?

To evaluate results from different models and feature sets, we followed multiple approaches described in this and the following subsections. First, to get a high-level understanding of the predictions made by our models, we performed exploratory GO enrichment analyses of genes predicted with high confidence to be involved in RNA methylation. Here, we defined as high confidence all genes in the top 1% of the probability distribution for Class 1. For the GB ensemble trained on the full feature set, this comprised the top 269 predictions with an average probability score greater than 0.83. For the SVM models trained on the reduced feature set, 268 genes with a probability of 0.84 or higher were selected.

The top 50 enriched terms for GB and SVM models are shown in Figures 4A and B, respectively. Both model ensembles, independently of the dataset they derived from, yielded predictions enriched in GO terms associated with RNA biogenesis, localization, transport and processing. Note that top enrichment results for GB included additionally terms associated with DNA and protein methylation processes (Figure 4A). This may point to either a lack of specificity of the models with regards to the modification substrate, or a close functional link between RNA and other methylation pathways. Overall, the GO analyses provided a good qualitative control for model performance. The rationale here is that although we did not recover enrichment in the biological term “RNA methylation” *per se* (given that the models predict “novel” genes), features closely associated with the term should figure among the top GO results.

**Figure 4.**
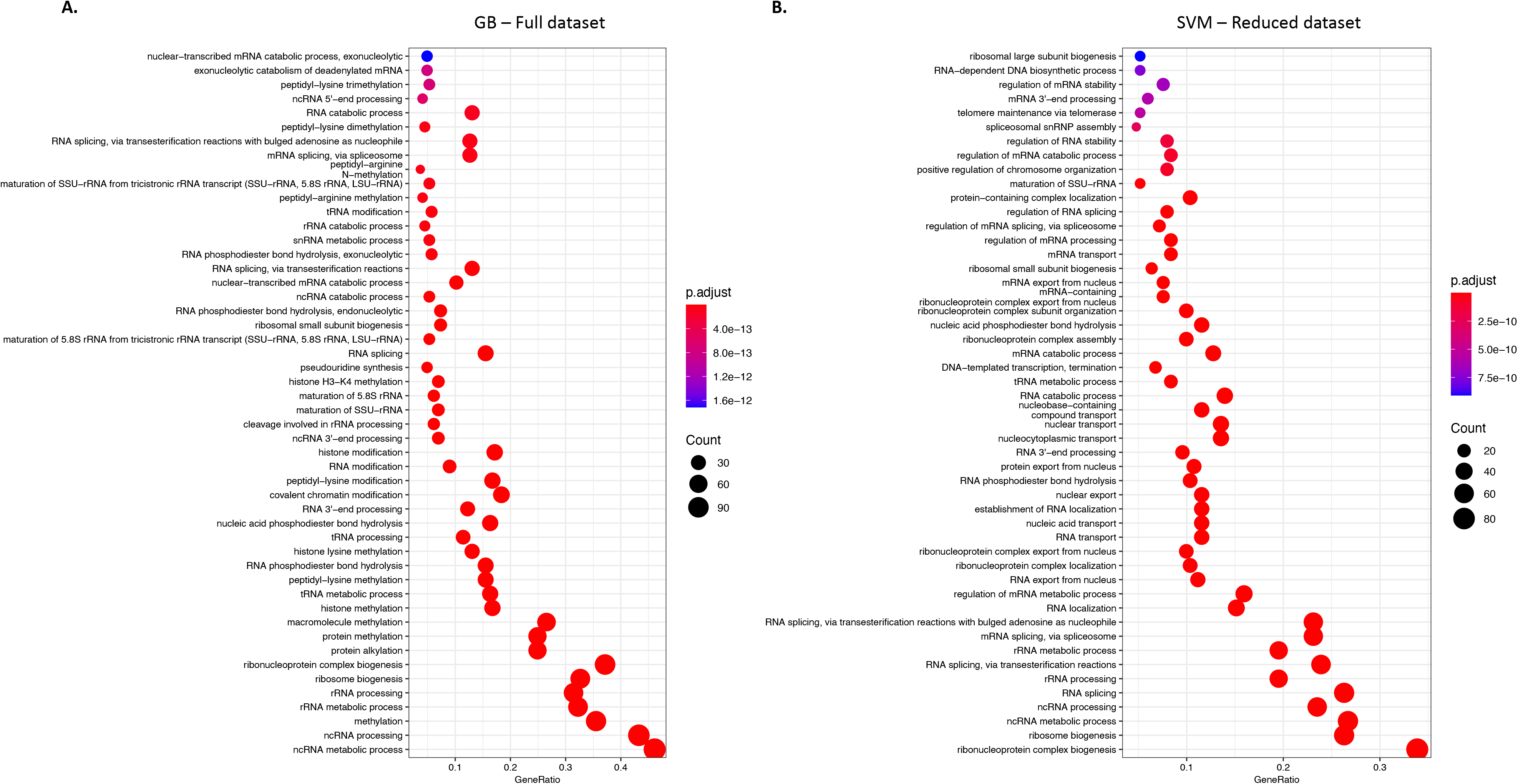
Functional enrichment analyses of high-confidence predictions. GO enrichment analysis of all genes in the top 1% of the probability distribution for Class 1 based on **A.** GB models, full feature set and **B.** SVM models, reduced feature set. Top enriched terms include functions such as RNA biogenesis, localization, transport, and processing. For GB predictions, additional functions were associated with DNA and protein methylation processes.

#### Do the models agree?

Our second analysis aimed to assess the degree of concordance between predictive models trained on the full and reduced feature sets. Figure 5 shows the predicted probability scores of each gene being assigned to Class 1, based on GB models derived from the full feature set versus the average probability obtained by the SVM models trained on the reduced feature set. Overall, the two ensembles yielded very similar predictions, as exemplified by the strong correlation between predicted probability scores (r = 0.872, P < 2.2e-16). Yet, for certain genes we observed a high degree of discordance between the GB/full and SVM/reduced models.

**Figure 5.**
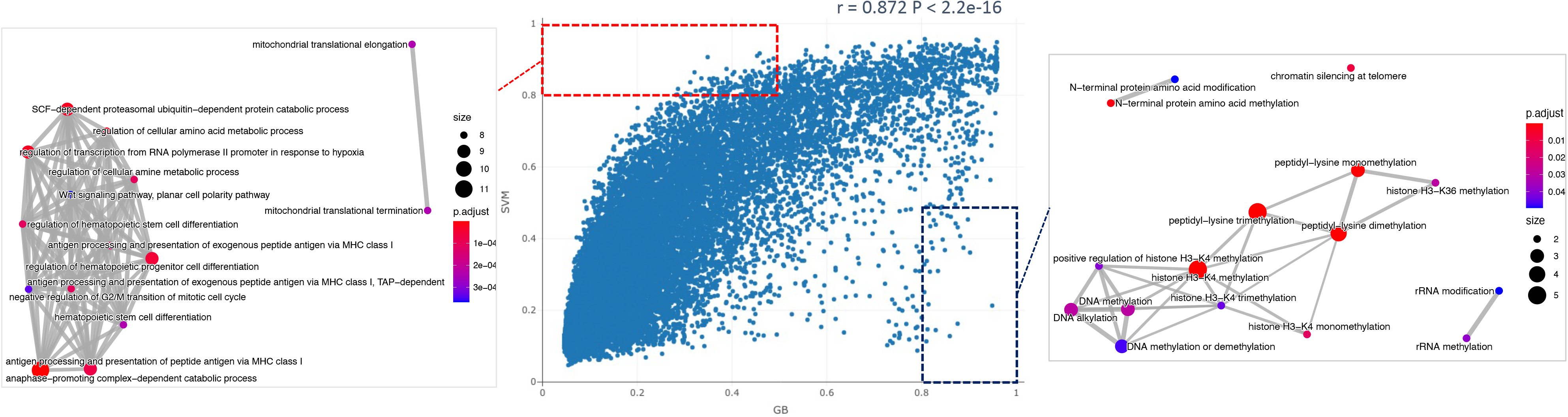
Concordance between predictive models. Middle panel: Scatterplot of the predicted probability score of each gene being assigned to Class 1, based on GB models trained on the full feature set versus SVM models trained on the reduced feature set. Side panels: Top 15 enriched GO terms associated with genes assigned to Class 1 with a probability greater than 0.8 by one ensemble only (right: SVM models only; left: GB models only). Enriched terms are represented as a network with edges connecting overlapping gene sets.

To further explore these discrepancies, we examined genes predicted to associate with RNA methylation pathways with a probability greater than 0.8 by one ensemble, but that were assigned to the negative class (P < 0.5) by the other ensemble. GO analysis of RNA methylation genes only predicted by SVM showed enrichment in the functions of anaphase-promoting complex-dependent catabolic process (P = 2.60E-07), antigen processing and presentation of peptide antigen via MHC class I (P = 7.69E-05), and mitochondrial translational elongation (P = 2.43E-04) among others (Figure 5). Given that gene expression constituted the most informative feature type for classifiers trained on the reduced feature set, it is likely that genes participating in the aforementioned processes exhibit highly similar expression profiles to RNA methylation genes - at least according to transcriptomic resources used here for learning.

On the opposite end of the distribution, considering genes recovered with a high probability score by GB models only, our analyses found significant enrichment in DNA, histone and protein methylation processes, as well as other RNA modification pathways (P < 0.05, Figure 5). This may represent a modelling artifact, i.e., predictions erroneously assigned to Class 1, that could be caused by the hierarchical nature of GO terms (e.g., “methylation” being the parent term of both “RNA methylation” and “DNA methylation” processes). An alternative interpretation is that our models capture a functional link between modification pathways operating at different substrates.

#### In silico validation of gene predictions

Of all classifiers, GB models that were trained on the full feature set showed the best performance based both on cross-validation and hold-out test datasets. We thus selected the top hundred genes predicted by the GB models to associate with RNA methylation pathways as candidates for further validation (Table 5). To evaluate these predictions with respect to previously known RNA methylation genes, we first performed a hierarchical clustering analysis of predicted plus positive (Class 1) genes based on the machine learning data used here (Figure 6). As anticipated, known and predicted genes were well clustered together, with no evident split between known and predicted RNA methylation genes.

**Figure 6.**
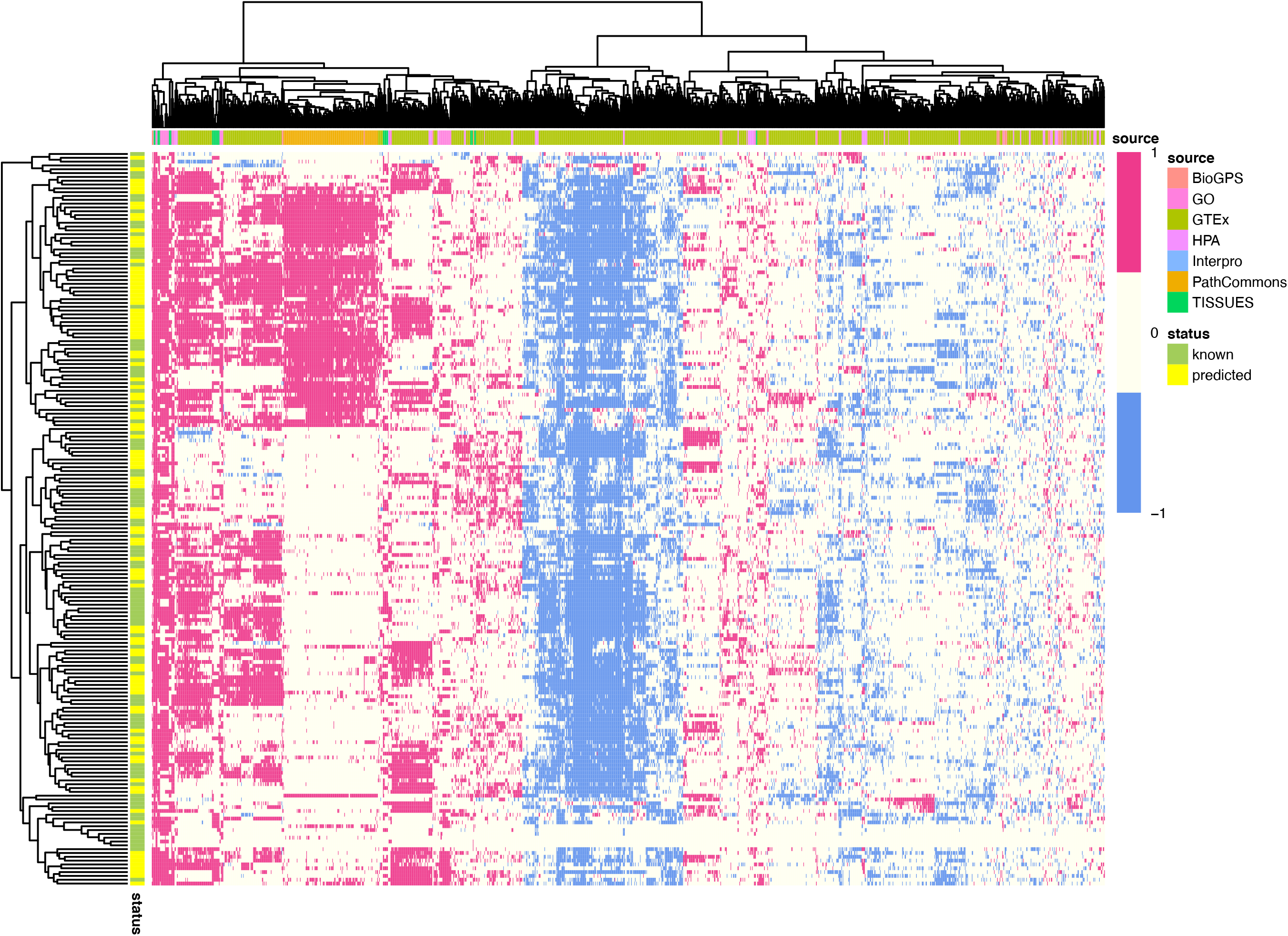
Heatmap of predicted and known RNA methylation genes. Hierarchical clustering analysis of predicted plus positive genes shows no evident split between predictions (yellow) and known RNA methylation genes (green). Features (columns) used for machine learning are shown in different colours based on the data source.

**Table 5.**
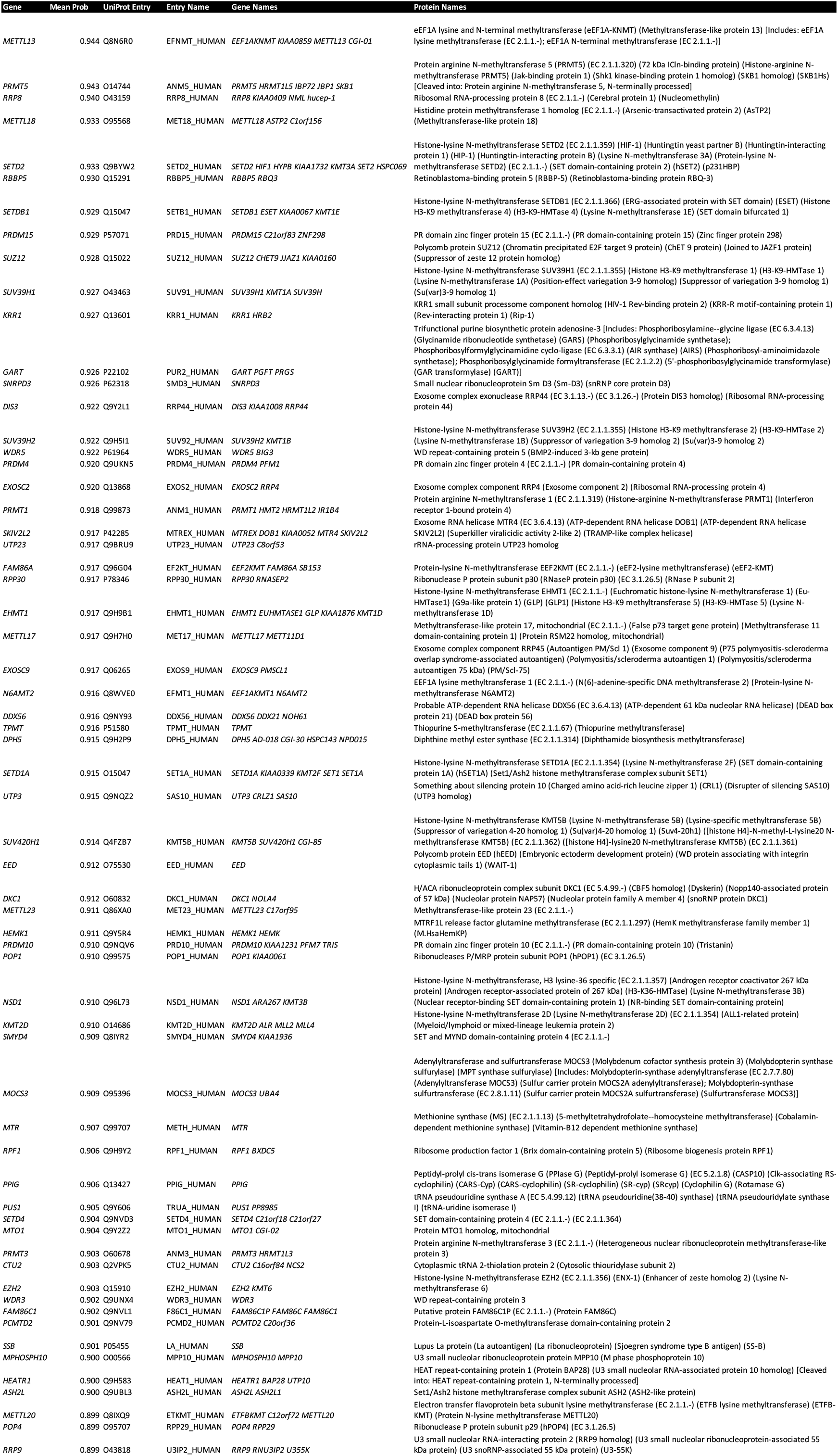
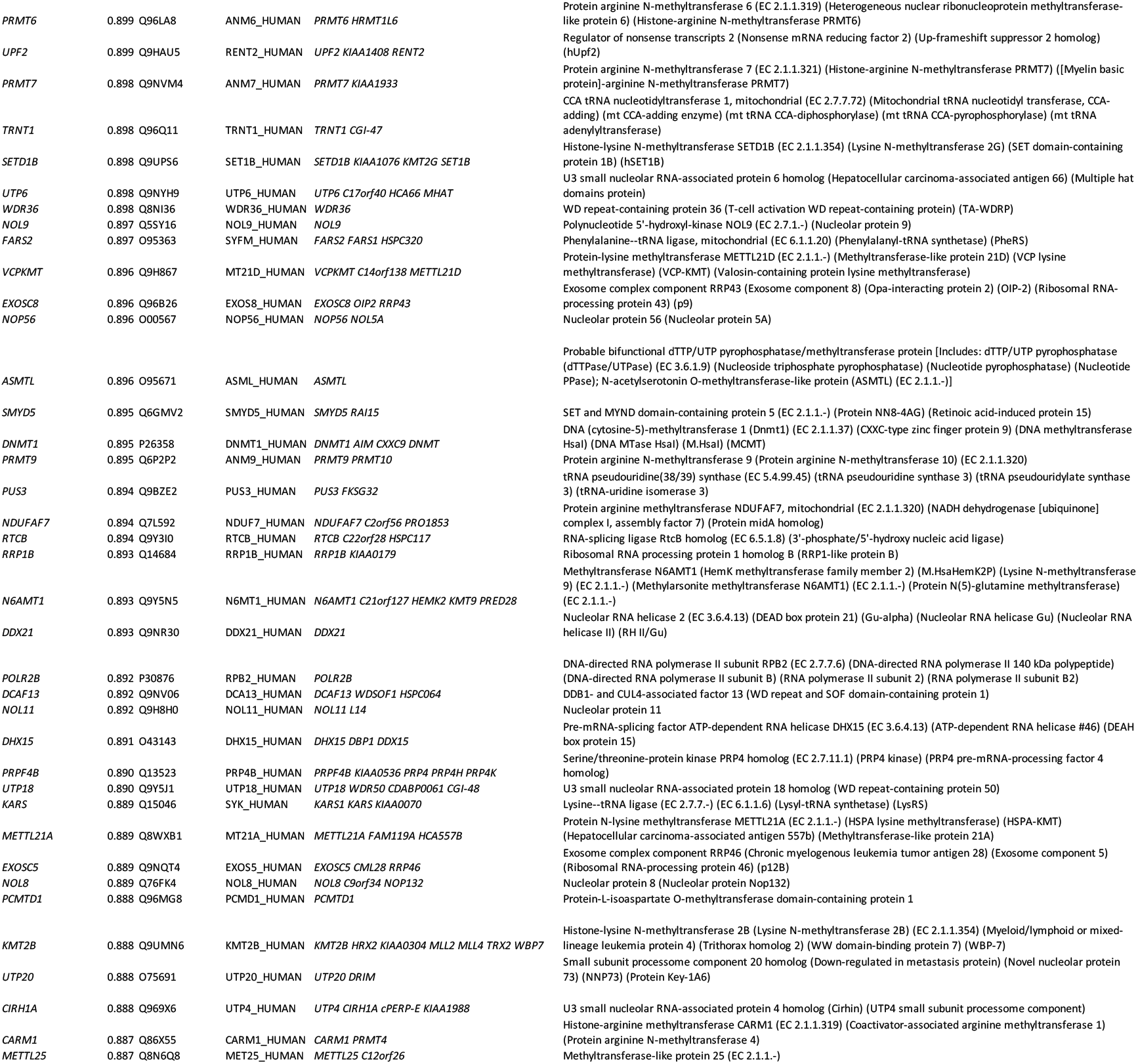
Top 100 gene predictions based on the GB model ensemble of the full feature set.

Second, we interrogated the STRING database^21^ for independent Protein-Protein Interaction (PPI) information on known RNA methylation genes and other genes of the human genome. We built a PPI network based on interactions with a confidence score of 400 or above, and performed Random Walks starting from proteins known to mediate methylation of RNAs (Class 1). This allowed us to weigh all other proteins in the network and rank them by their importance relative to our positive gene set. To evaluate whether genes predicted by our models were highly ranked among important interactors, we performed Gene Set Enrichment Analysis (GSEA) using the PageRank score as an input. We obtained a strong positive enrichment (NES = 1.605, P = 0.0001) for the model predictions (Table 6), corroborating their close functional association with RNA methylation pathways based on independent PPI evidence (Figure 7).

**Figure 7.**
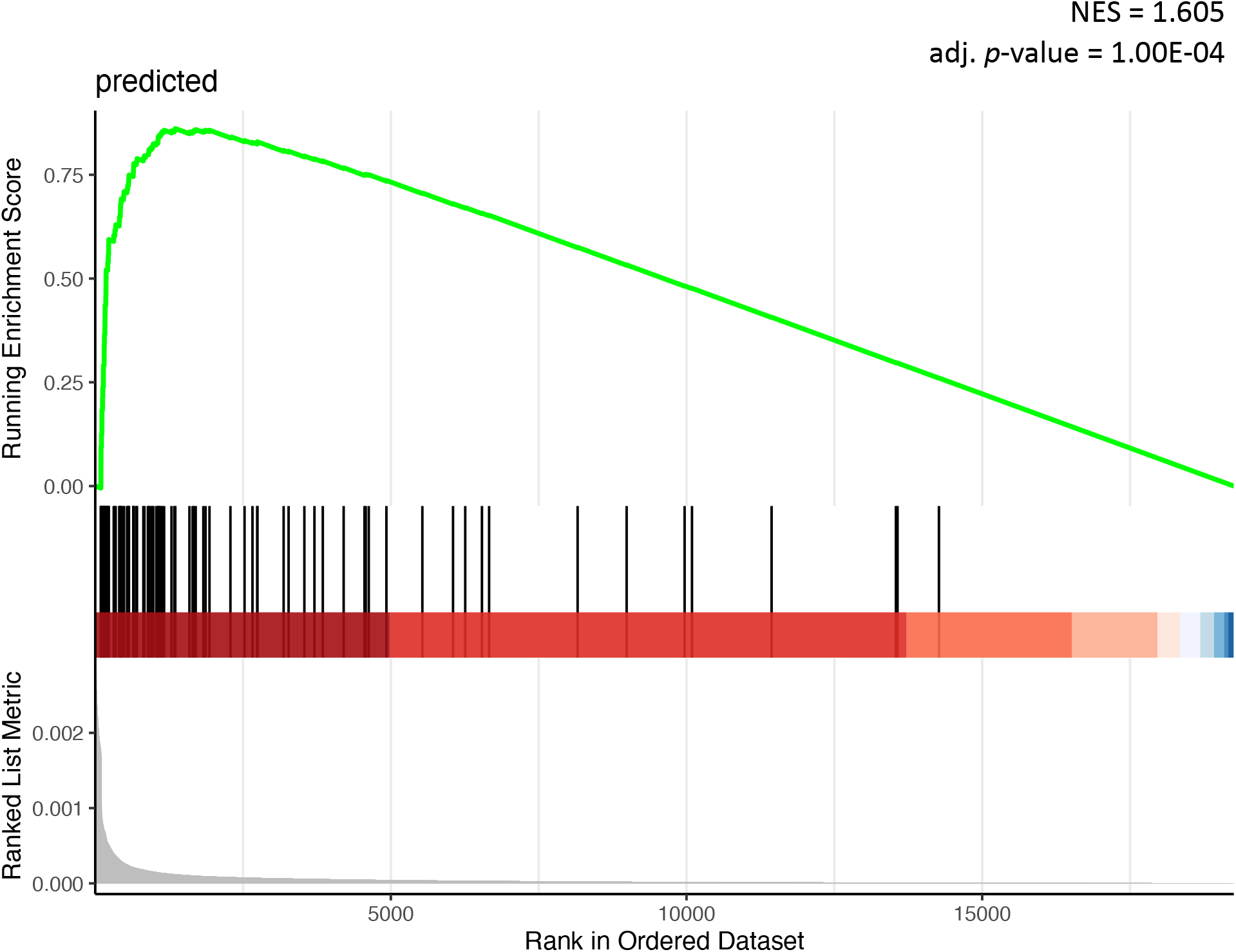
GSEA analysis of model predictions based on PageRank score. Personalised PageRank score of all human genes was computed using PPI data from STRING, starting from previously known RNA methylation genes. A strong positive enrichment (NES = 1.605, P = 0.0001) was obtained for model predictions, corroborating a close functional association with RNA methylation pathways.

**Table 6.**
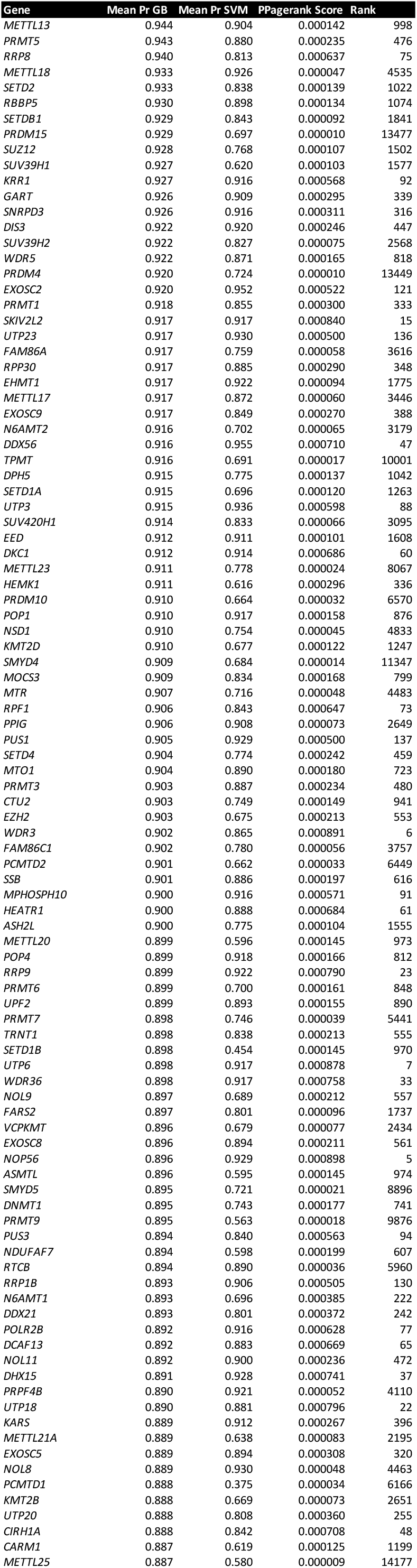
Personalised PageRank score of top 100 model predictions based on PPI data (source: STRING).

### Insights into the role of new predictions

To gain functional insights into the role of newly predicted genes with regards to previously annotated RNA methyltransferases and associated proteins, we interrogated the STRING database for available PPI data connecting our model predictions to known RNA methylation genes. Our search unravelled a dense network of interactions (Figure 8A), comprising 2,450 edges (confidence ≥ 400). To further dissect these PPI data and identify subgroups of proteins associated with specific pathways, we employed the Louvain method of community detection22. We identified six communities in total (Figure 8B), which we annotated using a large collection of functional annotation resources^23^.

**Figure 8.**
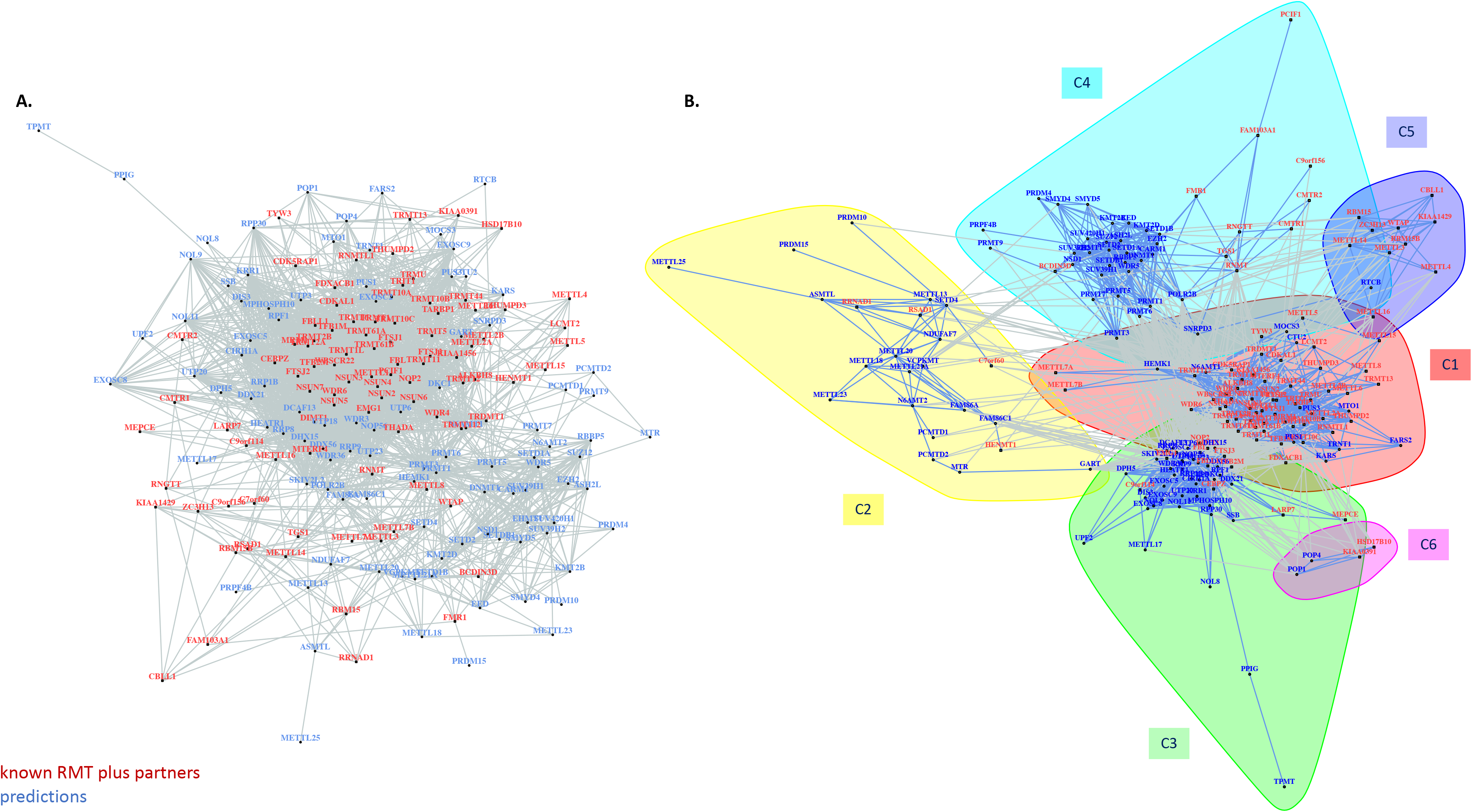
PPI network of known and predicted genes involved in RNA pathways. **A.** Network based on available PPI data connecting newly predicted genes with previously annotated RNA methyltransferases and associated proteins. **B.** Subgroups of proteins associated with specific pathways, as inferred using the Louvain method of community detection.

Community 1 (C1, Figure 8B) groups most RNA methylation genes from the positive set, together with 10 model predictions: *CTU2*, *FARS2*, *HEMK1*, *KARS*, *MOCS3*, *MTO1*, *N6AMT1*, *PUS1*, *PUS3* and *TRNT1*. Functional analysis of community members showed that proteins comprising this sub-network are significantly enriched in the functions of tRNA modification (GO:0006400, P = 5.09E-70), tRNA methylation (GO:0030488, P = 6.31E-66), and tRNA processing (Reactome R-HSA-72306, P = 4.10E-45). Indeed, four predictions in the cluster, CTU2, MOCS3, PUS1 and PUS3, are RNA modifying enzymes mediating tRNA modifications. CTU2 and MOCS3 are involved in 2-thiolation of mcm^5^S^2^U at wobble positions of tRNAs, whereas PUS1 and PUS3 belong to the tRNA pseudouridine synthase TruA family and mediate the formation of pseudouridine at positions 27/28 and 38/39 of certain tRNAs, respectively^13^. Among other members of the same community, the gene *TRNT1* encodes the mitochondrial CCA tRNA nucleotidyltransferase 1 responsible for the addition of the conserved 3’-CCA sequence to tRNAs. It has been previously reported that the presence of the 3’-CCA tail on tRNA is required for target recognition by the tRNA methyltransferase NSUN6^24^, which could underlie the functional link of TRNT1 with RNA methylation genes in our analyses.

Likewise, two aminoacyl-tRNA synthetases, FARS2 and KARS, were also predicted to be closely associated with RNA methylation pathways and were part of Community 1. FARS2 is a mitochondrial Phenylalanine-tRNA ligase, responsible for the charging of tRNA(Phe) with phenylalanine. *KARS* encodes a Lysin-tRNA ligase. Although, we have not found any orthogonal evidence linking FARS2 to RNA methylation, KARS has been previously inferred to physically interact with the RNA methyltransferase TRMT1, based on co-fractionation data (source BioGRID^25^).

The same sub-network also included two HemK methyltransferases, HEMK1 and N6AMT1. The former is a N5-glutamine methyltransferase responsible for the methylation of the glutamine residue in the GGQ motif of the mitochondrial translation release factor MTRF1L^26^. N6AMT1 methylates the eukaryotic translation termination factor 1 (eRF1) on Gln-185. Notably, it has been reported that N6AMT1 forms the catalytic subunit of a heterodimer with the RNA methyltransferase TRMT112^27^, suggestive of a functional interplay between RNA methylation and post-translational modifications of translation factors.

Our models also predicted that *MTO1* is a gene functionally associated with RNA methylation pathways. Previous studies have shown that *MTO1* encodes for a mitochondrial protein which is indeed involved in the 5-carboxymethylaminomethyl modification (mnm^5^s^2^U34) of the wobble uridine base in mitochondrial tRNAs, with a crucial role in translation fidelity^28^.

Community 2 (C2, Figure 8B) consists mainly of newly predicted genes, associated with four genes from the positive set: *C7orf60*, *HENMT1*, *RRNAD1* and *RSAD1*. The gene *C7orf60* or *BMT2* encodes a probable S-adenosyl-L-methionine-dependent methyltransferase. Recent studies have suggested that BMT2 (also known as SAMTOR) acts as an inhibitor of mTOR complex 1 (mTORC1) signalling in human, a SAM sensor signalling methionine sufficiency^29^. In yeast, BMT2 is responsible for the m^1^A2142 modification of 25S rRNA^30^. Two other methyltransferase genes in the same cluster were *RRNAD1* and *HENMT1*. The former encodes for ribosomal RNA adenine dimethylase domain containing 1, but little is known about its function. HENMT1 is a small RNA methyltransferase that adds a 2’-O-methyl group at the 3’-end of piRNAs, contributing to the maintenance of Transposable Element (TE) repression in adult germ cells^31^. Functional annotation of this community indicated an enrichment in peptidyl-lysine methylation function (GO:0018022, P = 1.92E-06), albeit this was based on only four proteins out the 23 forming this cluster (SETD4, VCPKMT, METTL21A, and METTL18). Among members of this community, we identified proteins with a role in methylation of other substrates. For example, FAM86A catalyses the trimethylation of the elongation factor 2 (eEF2) at Lys-525^32^. METTL13 is also a methyltransferase responsible for the dual post-translational methylation of the elongation factor 1-alpha (eEF1A) at two positions (Gly-2 and Lys-55), modulating mRNA translation in a codon-specific manner^33^. Both genes are involved in modifying translation elongation factor residues, same as N6AMT1 mentioned above. Our results hence suggest that post-translational modifications of translation factors and epitranscriptomic changes on RNAs could be interconnected in modulating translational efficiency.

Community 3 (C3, Figure 8B) comprises 48 protein members, of which 10 are part of our positive set and 38 were predicted by the models. Overall, we found a strong enrichment for functional terms linked to ncRNA processing (GO:0034470, P = 6.79E-40) and rRNA processing (R-HSA-72312, P = 1.03E-39). For example, among Community 3 members, our predictions include five genes encoding for members of the nuclear RNA exosome, *DIS3*, *EXOSC2*, *EXOSC5*, *EXOSC8* and *EXOSC9*. The exosome is known to participate in a wide variety of cellular RNA processing and degradation events preventing nuclear export and/or translation of aberrant RNAs. Exosome function is thus likely to be interlinked with epitranscriptomic marks on RNAs.

We also identified a sub-cluster within the community connecting DIMT1, EMG1, FBL and NOP2 with 15 proteins predicted by our models. All members of the sub-cluster are RNA-binding proteins involved in rRNA modification in the nucleus (R-HSA-6790901, P = 5.44E-36). *EMG1* encodes for an RNA methyltransferase that methylates pseudouridine at position 1248 in 18S rRNA^34^. Pathway annotation data further suggest that EMG1 together with eight new predictions (CIRH1A, DCAF13, HEATR1, NOL11, UTP3, UTP6, UTP20 and WDR3) are required in pre-18S rRNA processing and ribosome biogenesis. Of these, the *NOL11* gene encodes a nucleolar protein contributing to pre-rRNA transcription and processing^35^. Partial evidence furthermore suggests that NOL11 interacts with the rRNA 2’-O-methyltransferase fibrillarin, FBL, which is involved in pre-rRNA processing by catalysing the site-specific 2’-hydroxyl methylation of pre-ribosomal RNAs^35^. FBL together with RRP9 and NOP56 are part of the box C/D RNP complex catalysing the ribose-2’-O-methylation of target RNAs.

Finally, three novel gene predictions within this community, *DPH5*, *TPMT* and *RRP8*, were previously reported to have SAM-dependent methyltransferase activity. *DPH5* is coding for a methyltransferase that catalyses the tri-methylation of the eEF2 as part of the diphthamide biosynthesis pathway, whereas *TPMT* encodes an enzyme that metabolizes thiopurine drugs. We cannot rule out that these may be false positives cases, i.e., erroneous predictions that stem from the presence of the SAM-binding domain in the protein. Yet genes mediating post-translational modifications were repeatedly classified as components of RNA methylation pathways by our machine learning models (e.g., *FAM86A* in Community 2). A noteworthy case is RRP8, which in human is reported to bind to H3K9me2 and to probably act as a methyltransferase, yet studies in yeast have shown that the RRP8 homologue is responsible for installing m1A in the peptidyl transfer centre of the ribosome (m^1^A645 in 25S)^36^.

Community 4 (C4, Figure 8B) constitutes a large cluster of 42 proteins. Functional analysis of the group indicates that most community members are chromatin modifying enzymes (R-HSA-3247509, P = 8.74E-29), or are associated in general with chromatin organization (R-HSA-4839726, P = 8.74E-29) and histone modification (WP2369, P = 1.08E-23). Previously known RNA methylation genes in this community were mainly involved in RNA-capping pathways, e.g., *RNMT*, *CMTR1*, *CMTR2*, *FAM103A1*, *TGS1* and *RNGTT*. Recent studies have suggested that there is indeed extensive crosstalk between RNA modifications and epigenetic mechanisms of gene regulation^7,37,38^.

Community 5 (C5) and Community 6 (C6) encompass fewer members than the other communities. Community 5 consists of 10 proteins creating a small sub-network of RNA methyltransferases and partner proteins involved in RNA methylation (GO:0001510, P = 1.91E-17) and mRNA methylation, in particular (GO:0080009, P = 6.26E-16). Notably, this community captures proteins involved in the m6A pathway, including the m^6^A writer complex of METTL3-METTL14 with co-factor WTAP, METTL16 and ZC3H13, as well as the m^6^Am writer METTL4^39^. Community 6 is the smallest of all communities with only four protein members, two previously annotated RNA methylation genes, *HSD17B10* and *KIAA0391*, and two predicted genes *POP1* and *POP4*. Functional analysis suggests that all four proteins contribute to tRNA processing (R-HSA-72306, P = 5.97E-09) and three of them are involved in tRNA 5’-end processing (GO:0099116, P = 5.32E-08). The *HSD17B10* gene encodes the 3-hydroxyacyl-CoA dehydrogenase type-2, which is involved in mitochondrial fatty acid beta-oxidation. *HSD17B10* is involved in tRNA processing as it also forms a subcomplex of the mitochondrial ribonuclease P together with TRMT10C/MRPP1^40^. This subcomplex, named MRPP1-MRPP2, catalyses the formation of N1-methylguanine and N1-methyladenine at position 9 (m^1^G9 and m^1^A9, respectively) in tRNAs. *KIAA0391*, also known as *PRORP*, encodes a catalytic ribonuclease component of mitochondrial ribonuclease P. It appears that POP1 and POP2 are also components of ribonuclease P and contribute to tRNA maturation via 5’-end cleavage.

### Potential drawbacks

Our machine learning models and analyses have provided a wealth of new information on putative gene networks underpinning RNA methylation in human. However, it is worth noting the limitations of our approach. First, because only few writer enzymes are to date known to deposit methyl-marks on RNA6, we started from a very limited number of positive (and by consequence negative) samples to use for machine learning. Even though model performance based on test data was good, the small sample sizes may have hampered how well our models generalise. In addition, our models overpredicted genes associated with RNA methylation pathways, as a large number of genes obtained a high probability score for Class 1. This is because we followed a modelling approach using balanced positive and negative classes to optimise model performance.

Second, it is uncertain whether employing previous knowledge from functional annotations may have biased model predictions. We addressed this caveat to an extent by using a reduced feature set without annotation features, such as GO terms. When looking at predictions based on models trained on this dataset, we identified genes previously known to be involved in cell differentiation, G2/M cell cycle, antigen presentation and mitochondrial translation (P < 0.05, Figure 5). Even based on this unbiased set of classifiers, machine learning models point to a recurrent theme of this study: that RNA methylation is functionally interconnected to a range of other core cellular functions. For example, we repeatedly found genes encoding protein methyltransferases among the top model predictions. The key question here is whether these genes represent false positives, spurred by the hierarchical structure of GO terms or the shared SAM binding domain. These ambiguous predictions should be interpreted with caution, although multiple lines of evidence suggest that this could well be a biologically meaningful result echoing the crosstalk between DNA, RNA and post-transcriptional modification processes.

## CONCLUSIONS

RNA methylation is a key modulator of transcript stability, splicing and translation efficiency, playing a critical role in cellular homeostasis and disease^4^. Yet, its molecular underpinnings remain to date poorly understood^11^. Here, we aimed to gain novel insights into genes associated with RNA methylation pathways in human using machine learning approaches. Specifically, we analysed available transcriptomic, proteomic, structural and protein-protein interaction data in a supervised machine learning framework.

Our machine learning models showed very good performance on unseen test data, reaching high accuracy (91%), precision (90%) and recall (92%). *A priori* gene knowledge (e.g., GO annotations) together with expression data constituted the most informative data types in predictive modelling. Notably, in certain tissues, such as blood, heart, pancreas and brain, genes mediating RNA methylation seemed to show an up- or down-regulated expression profile.

Using independent PPI data, we orthogonally validated top model predictions by corroborating close functional links to previously known RNA methylation genes. Community detection delineated six molecular subnetworks, with distinct roles in tRNA processing (C1, C6), rRNA processing (C3), mRNA methylation (C5), but also protein (C2) and chromatin modifications (C4). Network analyses suggested that deposition of methyl marks on tRNAs is co-orchestrated with other modification processes, such as 2-thiolation and pseudouridine formation. Similarly, rRNA methyltransferases appeared functionally linked to several genes involved in rRNA processing and ribosomal biogenesis. Intriguingly, RNA-capping enzymes were clustered with chromatin modifiers, raising the hypothesis of a crosstalk between the two processes. Our results further indicate that post-translational modifications of translation factors and epitranscriptomic changes on RNAs are intertwined in modulating translational efficiency. Overall, our study exemplifies how access to omics datasets joined by machine learning methods can be used to infer molecular pathways and novel gene function.

## METHODS

### Dataset assembly and pre-processing

To assemble a machine learning dataset for predicting genes involved in RNA methylation process in the human genome, we first curated a list of previously known RNA methylation genes. For this, we performed searches in standard functional annotation resources, such as ExPASy ENZYME (https://enzyme.expasy.org/), InterPro (https://www.ebi.ac.uk/interpro/) and the GO Resource (http://geneontology.org/), in conjunction with a comprehensive literature review for annotated RNA methyltransferases following up on the pioneering paper of Schapira^6^. This allowed us to identify 92 proteins involved – or putatively involved – in RNA methylation to use for machine learning modelling (Table 1).

To obtain informative features for classifying gene functions, we interrogated the Harmonizome database^15^. Harmonizome provides a large collection of the pre-processed datasets for genes and proteins, with ~72 million attributes (functional associations) from over 70 major online resources. We selected 15 one-hot-encoded datasets from four broad categories: (i) transcriptomics; (ii) proteomics; (iii) structural or functional annotations; and (iv) physical interactions (Table 2). In particular, from omics experiments, we sampled BioGPS^16^, GTEx^18^, HPA^19^ and TISSUES^20^ gene and protein expression profile data. From functional datasets, we considered GO annotations and InterPro structural domains. Finally, from physical interactions datasets, we selected KEGG and Reactome Pathways, as well as Hub Proteins and Pathway Commons. Collating these data yielded an initial matrix of 26,935 genes and 50,176 one-hot-encoded features (“full feature set”). In addition, we compiled a second dataset of reduced dimensionality, by excluding all 5,148 GO and InterPro annotation features (“reduced feature set”).

### Problem framing, model definition, training and evaluation

To estimate the probability of a gene being associated with RNA methylation, we used standard machine learning approaches for binary classification. We labelled the 92 previously known RNA methylation genes as positive samples (Class 1), and split them into two sets comprising: (i) 80% of the data for training and cross-validation (n=74) and (ii) 20% kept unseen for model testing (n=18). We considered the remaining genes of the human genome as negative samples (Class 0) and performed an analogous 80/20 split into training/cross-validation (n=21,476) and test sets (n=5,368). The underlying assumption here is that the vast majority of genes in the human genome serve other functions, thus the number of false negatives in the training data should be very small.

To produce balanced sets of training samples, and to later reduce the variance of our final models through averaging, negative genes kept for training (n=21,476) were further divided into sets of 74 – equal to the number of positive samples for training. We thus generated 290 training sets, where the positive class remained fixed and the negative class was represented by a random draw of an equal number of genes from the rest of the genome, sampling each gene once.

Starting with 290 training sets and our unprocessed Harmonizome data comprising 50,176 features, we next performed filtering to remove low-information features. We removed features with (i) zero values in more than 70% of the samples in each training set, or (ii) less than 16% variance in at least one training set. The selected features for each of the 290 training sets were then merged into a final list of features for model training and testing. We followed the exact same selection process for the reduced feature set as well.

We next considered five types of machine learning models for binary classification: Logistic Regression (LR), Gaussian Naïve Bayes (GNB), Support Vector Machine (SVM), Random Forest (RF) and Gradient Boosting (GB) models. We used grid search and 3-fold cross-validation on each training set for the SVM hyperparameter tuning of the kernel function (linear or RBF), cost parameter, and kernel bandwidth (RBF kernel only). For RF, we used grid search to determine the optimal number of trees in the forest, followed by a randomized search to select the best parameters for maximum number of features considered for splitting a node, maximum number of levels in each decision tree, minimum number of data points placed in a node before the node is split, and minimum number of data points allowed in a leaf node. Likewise, for the GB model, we performed grid search to optimise the learning rate and number of trees in the forest, and subsequently performed a randomized search to tune the remaining decision tree parameters (see RF). We trained all five predictive models on each of the training sets from the full and reduced feature sets, respectively. The performance of all classifiers was estimated using 10-fold cross-validation, i.e., the dataset was split into 10 folds, of which nine were used for the training process and one for testing. The process was repeated ten times, and model performance was estimated using standard performance metrics: accuracy, precision, recall (sensitivity), F1 score and Area Under the Receiver Operating Characteristic Curve (AUROC), averaged across the ten repeats. Finally, we used GB feature ranking to determine the top 100 most informative features across the ensemble of training sets for the full and reduced feature sets, respectively.

### Final model testing on test dataset and genome-wide prediction

Once the best classifiers for the full and reduced datasets were selected based on cross-validation, we tested the performance of the model ensembles on unseen data. Analogous to the procedure described above for training data, we generated 298 testing datasets, by splitting the negative genes kept for testing into equal sets of 18 genes, and combining them with the 18 of positive samples previously retained. Each model from the classifier ensemble was evaluated on each of the test datasets using accuracy, precision, recall, F1 score and AUROC. Overall performance was calculated by averaging results of all models across test sets.

Likewise, the prediction probability of each human gene was calculated by averaging probability scores for Class 1 across all models of the best ensemble for the full and reduced feature sets, respectively. Most non-Class 1 genes (all except the test cases) were part of the negative samples in the training data of exactly one model in the ensemble; however, due to the high number of models (290) the effects of this on the final predictions is expected to be negligible.

All visualisations and meta-analyses were performed using the R software environment (v. 4.0.5)^41^. A heatmap of known and predicted RNA methylation genes across all features used for machine learning was generated using the R package pheatmap. Further *in silico* validation of model predictions was performed using GO enrichment analyses of predicted genes within the domain “Biological Process” using the package clusterProfiler^42^. Protein-Protein Interaction (PPI) data for human were obtained from STRING (v.11.0)^21^ and filtered to interactions with a combined score of 400 and above. All network analyses were performed using the igraph R package^43^. Functional annotation of PPI communities was performed using EnrichR^23^.

## ACKNOWLEDGEMENTS

The authors are thankful to Adrián Rodríguez Bazaga for his valuable input on the machine learning analyses, and Woochang Hwang for his feedback on the network analyses.

